# Multi-focal ultrasound neuromodulation to the dorsal anterior cingulate cortex disrupts behavioural and neural pain processing

**DOI:** 10.1101/2025.09.04.674195

**Authors:** Sophie Clarke, Samuel Mugglestone, Mathilde Lojkiewiez, Joshua Marquez, Nadège Bault, Elsa Fouragnan, Sam Hughes

## Abstract

Transcranial ultrasound stimulation (TUS) is a promising non-invasive technique for modulating deep brain regions involved in pain. TUS applied to the dorsal anterior cingulate cortex (dACC), a region implicated in chronic pain and established target for deep brain stimulation, has shown potential for reducing pain. This study aimed to investigate the neural mechanisms underlying TUS effects on pain in healthy participants using neuroimaging. Thirty-two participants underwent two double-blind, randomised TUS-fMRI sessions (active or sham). A tonic cold stimulus was applied during multifocal dACC-TUS and during fMRI and magnetic resonance spectroscopy (MRS) blocks. While no significant main effect of TUS on pain intensity was observed, active TUS showed a significantly greater reduction in pain ratings between 28- and 55-minutes post-stimulation, suggesting a delayed analgesic effect. Active TUS also disrupted the typical relationship between stimulus temperature and reported pain intensity, indicating altered sensory encoding. There was increased functional connectivity between the dACC and the supplementary motor area, pre-motor cortex, mid-ACC and the supramarginal gyrus, along with decreased coupling with the periaqueductal grey (PAG), and altered salience network connectivity. Overall, these findings suggest TUS to the dACC has multidimensional effects across behavioural and neural aspects of pain processing, supporting its potential therapeutic value.

## 1. Introduction

Transcranial ultrasound stimulation (TUS) enables precise, targeted neuromodulation of both cortical and deep brain regions, and it has been demonstrated across multiple studies that TUS can elicit behavioural changes and alter neuronal activity in humans, in both preclinical and clinical studies^1-4^. TUS is a particularly promising approach for pain neuromodulation given the involvement of deep cortical and sub-cortical regions in the pain network, such as the dorsal anterior cingulate cortex (dACC), insular cortex, thalamus or amygdala^5^. TUS protocols can be classified as either online; causing short-term effects during or immediately after stimulation, or offline; producing longer-lasting effects persisting beyond the stimulation period^1^. Previous studies have demonstrated effectiveness of both online and offline TUS in modulating pain, with online TUS applied to the dACC shown to reduce pain ratings to acute heat stimuli as well as alter autonomic responses and the amplitude of contact heat-evoked potentials, and offline TUS applied to the anterior thalamus shown to reduce thermal pain thresholds, both in healthy humans^6,7^. An early study in chronic pain patients showed offline TUS applied to the posterior frontal cortex resulted in significant improvements in pain ratings and mood 40 minutes after stimulation^8^. The delayed effects of offline TUS are particularly relevant in pain research, providing potential for translating these protocols into clinical interventions aimed at achieving long-lasting therapeutic benefits.

The dACC is a particularly relevant brain region for pain neuromodulation. It has a central role in pain processing, involved in both ascending and descending pathways and plays a role in emotional and cognitive processing^9^. Importantly, targeting the dACC using deep brain stimulation (DBS) has been shown to be effective in treating chronic pain conditions, with improvements in patient reported pain scores and quality of life^9-11^. In a cohort of chronic pain patients, TUS applied to the dACC has been shown to significantly reduce pain immediately after stimulation, with improvement sustained for up to 7 days post-treatment^12^.

The underlying neural basis for these improvements in reported pain following TUS to the dACC is not fully understood. This study aimed to further our understanding of the neural basis for altered pain processing following TUS to the dACC by using functional magnetic resonance imaging (fMRI) and magnetic resonance spectroscopy (MRS) techniques to explore functional connectivity changes and neurochemical changes respectively. TUS has been shown in several fMRI studies to illicit changes in functional connectivity with the brain region targeted, as well as larger scale changes in connectivity across brain networks^4,13-16^. MRS has also been previously employed to investigate neurochemical changes resulting from TUS, showing that TUS can result in altered GABA and glutamate (Glx) concentrations at the application site for some brain regions^4,17,18^, although one study also reported that GABA concentration was not altered following TUS to the dACC^4^. Multiple factors, such as the brain region, the current activation state or tissue composition, may influence TUS effects on neurochemical concentrations and further research is needed in this area. For the current study, we investigated TUS-induced changes in participant-reported pain ratings, functional connectivity and neurochemistry during exposure of the right hand to a tonic cold pain stimulus. TUS was applied to multi-sites of the dACC during tonic pain, to specifically elicit neuronal changes related to an active pain state in that brain region. TUS-induced effects were investigated at two timepoints, at 28 minutes post-TUS using fMRI measures of functional connectivity, and 55 minutes post-TUS using MRS measures of neurochemistry. We hypothesised that TUS would result in altered pain perception, accompanied by underlying changes in functional connectivity between brain regions known to be involved in pain processing as well as changes in neurochemical concentrations within the dACC.

## 2. Results

In total, 35 healthy participants were recruited to take part in the study, which investigated the effects of multi-focal active TUS vs sham TUS to the dACC on pain responses to a tonic cold stimulus (see study design summarised in Fig. 1A). Two participants were excluded after the first visit due to previously experiencing severe concussion (n=1), a contraindication for TUS, and due to an incidental finding in the first MRI (n=1). Therefore, 33 participants completed the study (mean age 26.3±10.2 years, range 21-66, 19 female, 14 male, sex and gender aligned by self-report). One dataset was excluded due to technical issues with the ultrasound, resulting in n=32 for the behavioural dataset, and three further datasets were excluded due to an artefact following scanner software upgrade, resulting in n=29 for the fMRI dataset.

**Figure 1.**
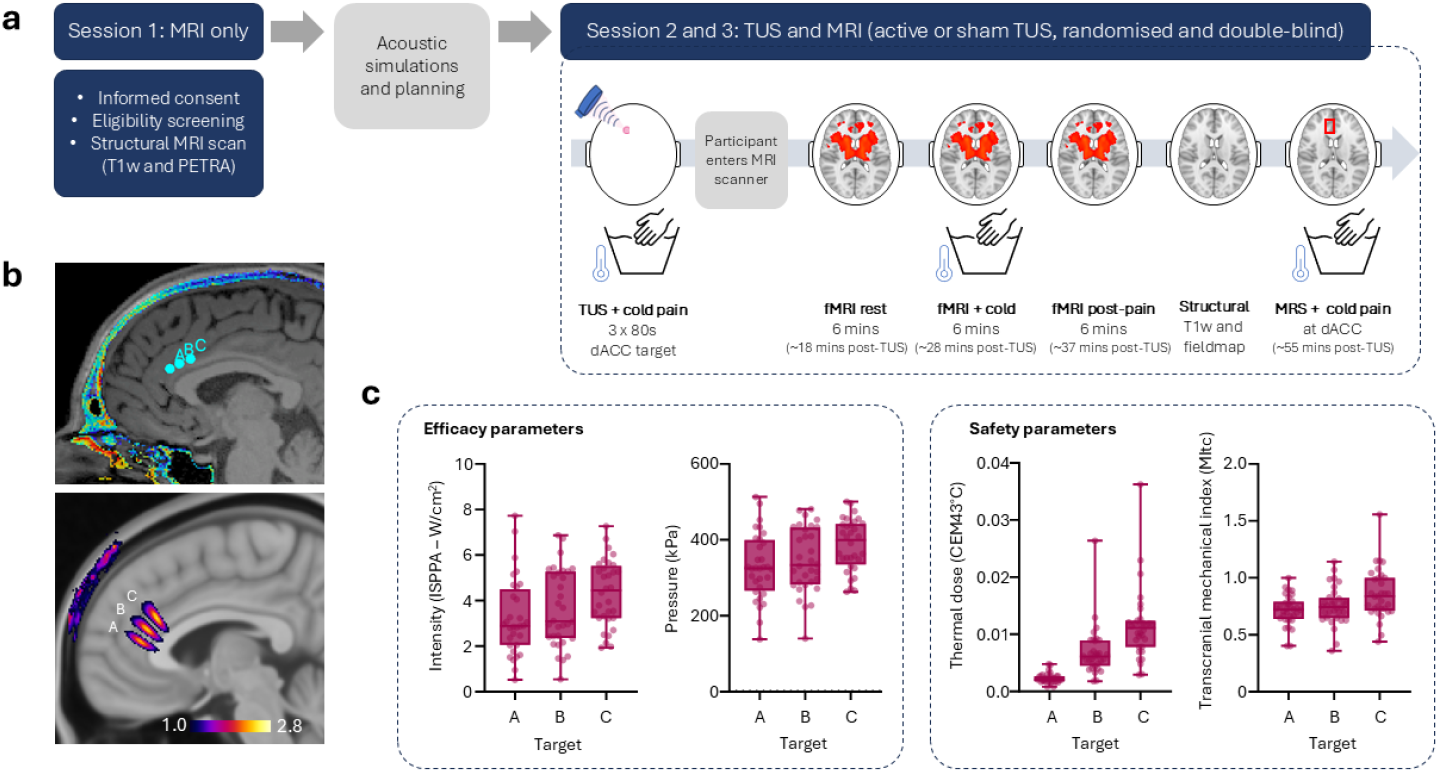
Study design and TUS post-stimulation simulation results (N=32) ***a*** Participants completed 3 sessions; in the first session, T1-weighted and PETRA scans were acquired for neuronavigation and acoustic planning, then at sessions 2 and 3 they received either active TUS to the dACC or sham followed by completing fMRI and MRS scans. **b** This multi-focal TUS intervention involved application of TUS to 3 target sites within the dACC, which are shown in the top panel labelled A, B and C. Average stimulation intensity (I_SPPA_) simulated using k-Plan software (BrainBox, Inc.) across all participants is shown in the bottom panel. **c** Efficacy and safety parameters from post-stimulation simulations shown for all participants for intensity (I_SPPA_) and pressure (kPa) in the dACC, thermal dose (CEM43°C) in soft tissues, and transcranial mechanical index (MItc) across the three targets.

### 2.1 TUS post-stimulation simulations for assessment of efficacy and safety

Targeted areas and average stimulation intensity (I_SPPA_) across all participants are shown in Fig. 1B. Detailed acoustic simulation parameters and outputs for all study participants can be found in Supplementary Materials [Table 1] and are summarised in Fig. 1C. Overall, there was good target engagement, with mean ISPPA in the dACC of 3.7±1.7w/cm^2^ across all participants. Importantly, transcranial Mechanical Index (MItc) remained below 1.9 and the Cumulative Equivalent Minutes at 43°C (CEM43), a metric reflecting both duration and intensity of heating relative to 43°C (the critical threshold for thermal cell damage), remained well below the safety threshold of 0.25^19^.

### 2.2 Behavioural results

Pain intensity ratings were given after approximately 6 mins tonic cold stimulus at three timepoints: T0 – during active or sham TUS to the dACC; T1 – during the resting state fMRI scan; and T2 – during the MRS scan. Linear mixed-effects modelling showed no significant main differences between active and sham TUS at T0, T1, or T2. The reduction in pain ratings from T1 to T2 was significantly greater in the active group compared to sham (Δ = –8.22, p = 0.049), potentially suggesting a delayed analgesic effect of TUS. This is shown in Fig. 2A.

**Figure 2.**
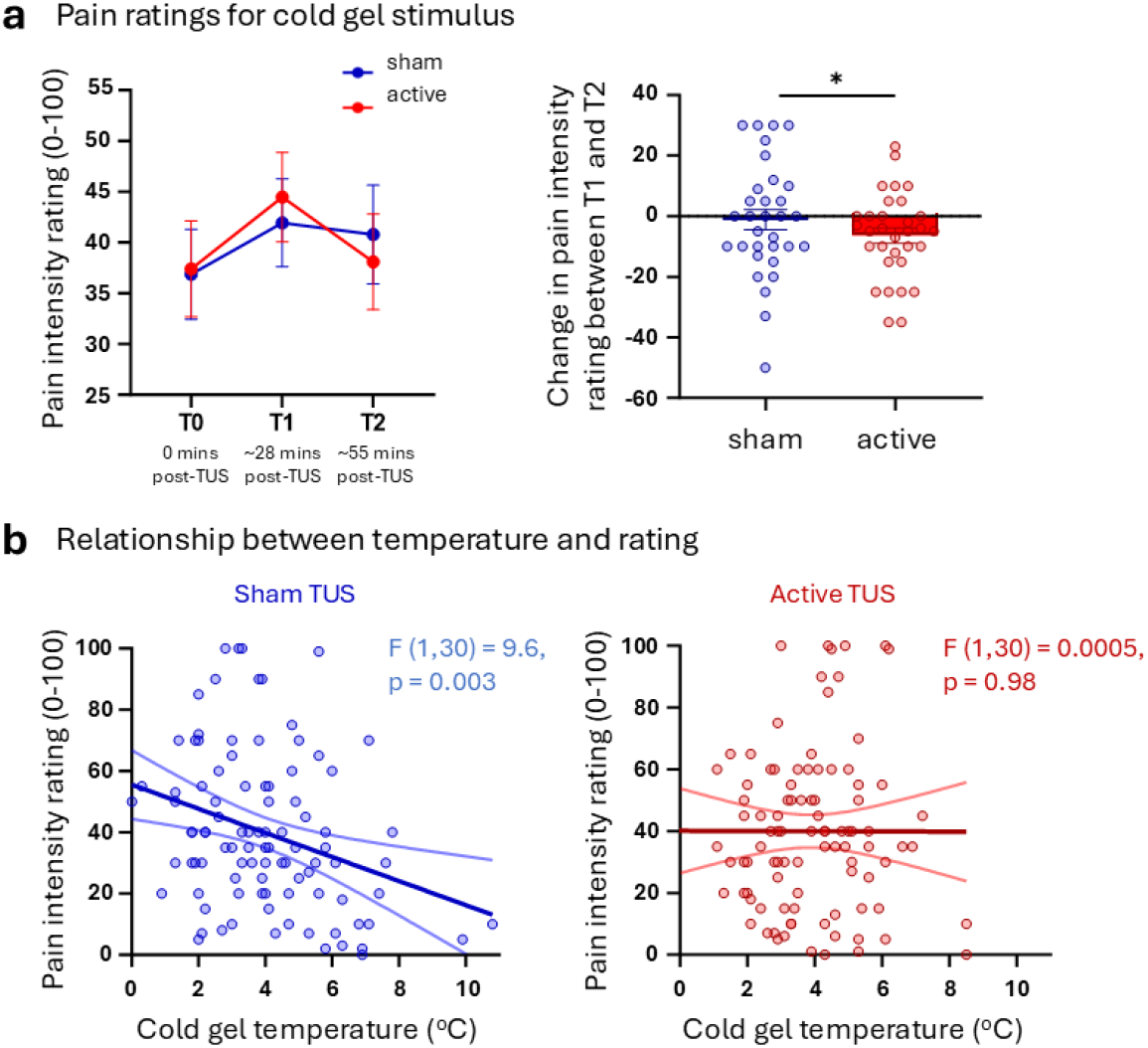
Pain intensity ratings to tonic cold stimulus (N=32) **a** Average pain intensity ratings for each of the three timepoints (left) and plot showing significant difference in the change in pain rating between T1 and T2 for active TUS compared to sham TUS (right). Error bars show the SEM. **b** Relationship between gel temperature and pain intensity ratings plotted for both sham TUS (left) and active TUS (right) showing the expected relationship between temperature and pain intensity is no longer present in the active TUS session.

A feature of the cold gelled-water stimulus used in this study was that there was a slight variation in temperature of the stimulus overall. This enabled investigation of temperature sensitivity across the pooled dataset. Simple linear regression was used to evaluate the relationship between temperature and pain intensity rating across all participants and all timepoints, for both the active TUS and sham TUS sessions. For sham TUS, there was a statistically significant relationship between temperature and pain intensity rating (R^2^ = 0.09, F (1,30) = 9.6, p = 0.003), whereas for active TUS sensitivity or intensity encoding to cold stimulus may have been altered as this expected relationship was no longer statistically significant (R^2^ = 4.8e-6, F (1,30) = 0.0005, p= 0.98). These results are shown in Fig. 2B.

### 2.3 Imaging results – seed-based functional connectivity and salience network connectivity

First, a seed-based connectivity approach was utilised to assess functional connectivity with the dACC target region. Whole-brain group average maps for mean activation during both the sham and active TUS conditions are shown in Fig. 3A, showing the connectivity profile of the dACC during the tonic cold stimulus which involves the thalamus, posterior cingulate cortex (PCC), caudate and putamen in both conditions. Differentially, there is connectivity between the dACC and the orbitofrontal cortex (OFC) in the sham condition, and with the anterior and posterior insular cortex, amygdala and supplementary motor area (SMA) in the active TUS condition. A whole brain, mixed effects analysis (Z>2.3, p<0.05) was conducted to compare active and sham TUS conditions, which showed there was increased connectivity between the dACC and pain-related brain regions, including the SMA, pre-motor cortex (PMC), mid-ACC and the supramarginal gyrus (SG), following active TUS compared to sham. This is shown in the bottom panel of Fig. 3A, with individual participant data plotted in Fig. 3B to illustrate this result showing that this reflects a change from negative dACC-SMA and dACC-PMC connectivity in the sham condition to positive connectivity in the active TUS condition. In addition to these two regions, functional connectivity coefficients for four further regions of interest, the thalamus, anterior insular cortex, dorsolateral prefrontal cortex (DLPFC) and periaqueductal grey (PAG), were also plotted (Fig. 3C). This illustrates the overall shift in the connectivity profile of the dACC, with the increased dACC-SMA and dACC-PMC connectivity accompanied by increased connectivity with the anterior insular and decreased connectivity with the PAG in the active TUS condition compared to sham, while dACC-thalamus and dACC-DLPFC connectivity was unchanged between conditions. These regions were selected as they are involved in pain responses and known to be connected to the dACC, with a focus on the left side as the pain stimulus was applied to the right hand.

**Figure 3.**
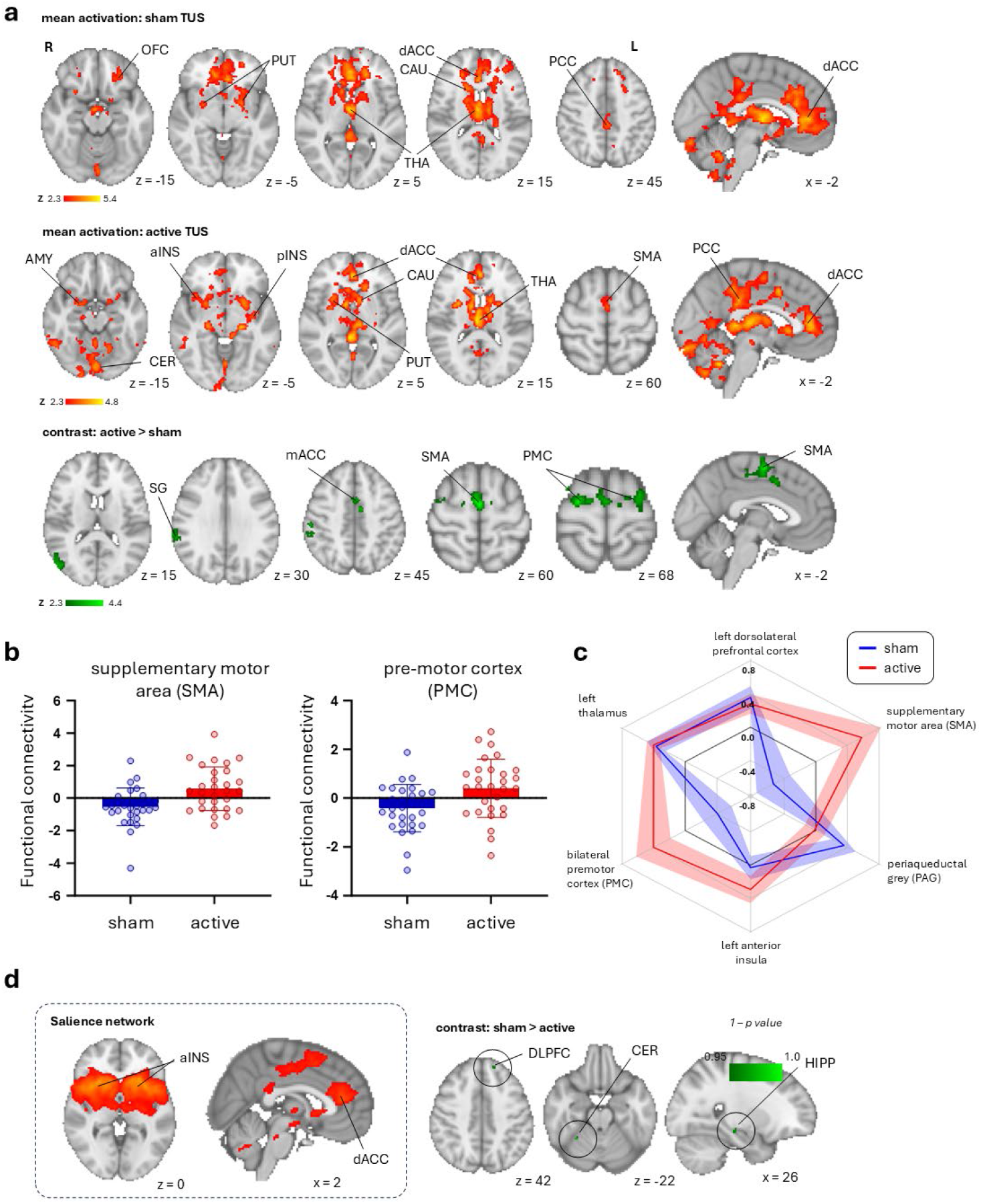
Seed-based functional connectivity and salience network connectivity results (N=29) **a** Whole-brain seed-based functional connectivity maps (mixed effects analysis, Z>2.3, p<0.05) with the dACC seed in the sham TUS condition (top row), the active TUS condition (middle row), and for the contrast active TUS condition > sham condition (bottom row). **b** Illustrative plots showing the altered connectivity with dACC seed region for the supplementary motor area (left), the pre-motor cortex (right). **c** Illustrative radar plot showing dACC connectivity with six different brain regions involved in pain processing, including the two regions that showed significantly altered connectivity in the whole-brain analysis, plus the left dorsolateral prefrontal cortex, the left thalamus, the left anterior insular cortex and the periaqueductal grey. **c** Salience network was identified using independent component analysis (ICA) carried out on the full dataset (shown in dotted inset), and subsequent group analysis comparing subject-specific network maps for active TUS and sham conditions showed altered salience network connectivity with the DLPFC, cerebellum and right hippocampus. MNI co-ordinates are shown for each image slice and error bars show the SEM.

Next, independent component analysis (ICA) and dual regression approaches were used to investigate changes in salience network connectivity – a well-defined brain network of which the dACC is a core node. Subject-specific salience network maps were compared for the active TUS and sham TUS conditions, and although there were no connectivity changes within the network itself, there was significantly decreased connectivity between the salience network and the left DLPFC, cerebellum and right hippocampus following active-TUS compared to sham-TUS. This is shown in Fig. 3D.

### 2.4 Metabolite concentration results

MRS data was collected at ∼55 minutes post-TUS to assess changes in gamma-aminobutyric acid (GABA) and glutamate (Glx) concentrations. Representative data from one participant is shown in Fig. 4A. Linear mixed effects modelling showed that there were no significant effects of session (active or sham TUS), sex or age for the concentration of GABA, Glx or the GABA/Glx ratio. The percentage change in GABA concentration between the sham and active TUS sessions is shown in Fig. 4B, illustrating there is no clear directional trend, and the GABA/Glx ratio is plotted for sham and active sessions in Fig.4C. There was a statistically significant relationship between the percentage change in GABA concentration and the change in slope for pain intensity ratings between the T1 and T2 timepoints (F (1,21) = 6.81, p = 0.02), shown in Fig. 4D. This relationship indicates that for participants where active TUS had a stronger excitatory effect on the dACC, resulting in a larger decrease in GABA concentration, there was a greater analgesic effect of TUS between T1 and T2 (based on a greater change in slope of pain ratings).

**Figure 4.**
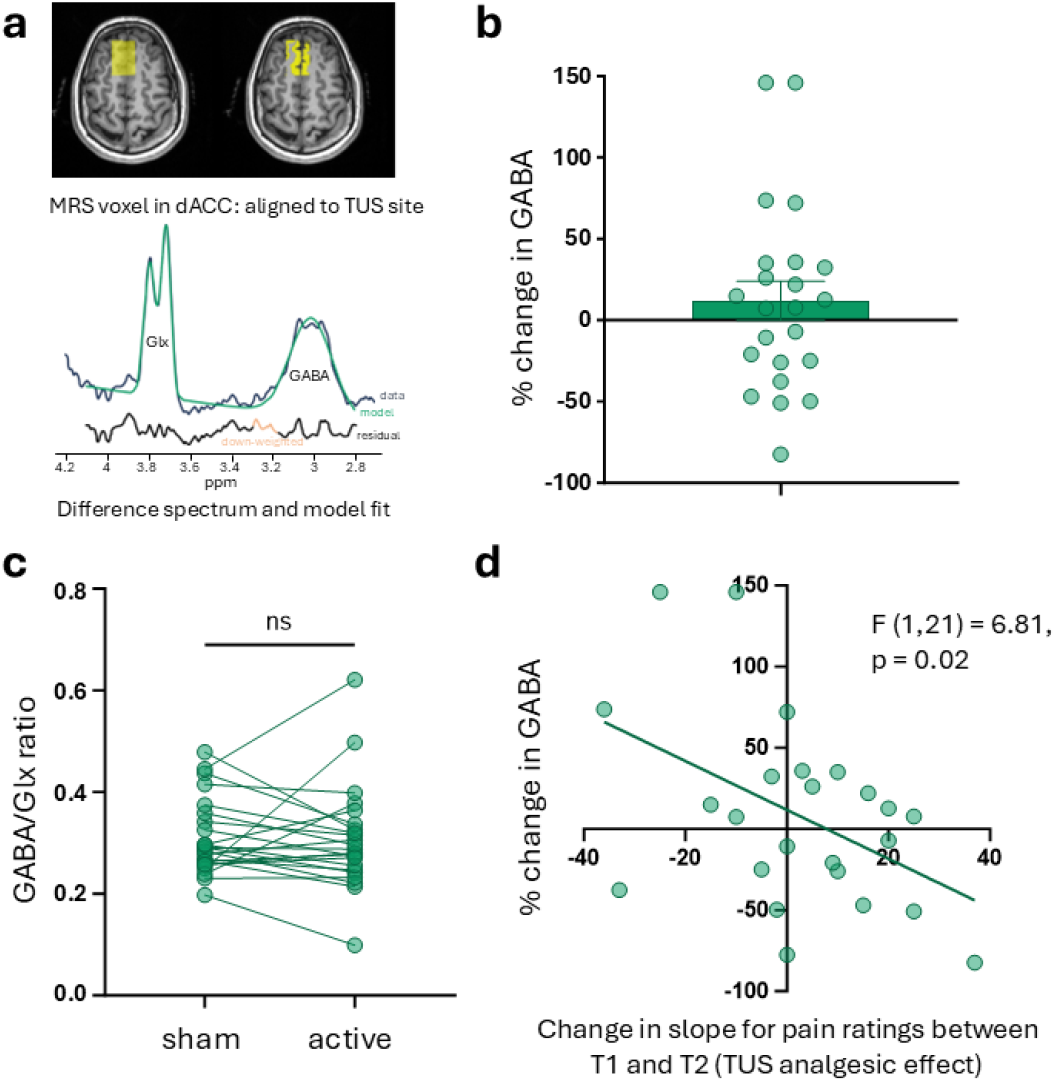
Metabolite concentration in the dACC (N=23) **a** Representative data from one participant showing the position of the MRS voxel over the dACC (top panel) and the difference spectrum and model fit (bottom panel), with the GABA spectrum in blue, model fit in red, and residuals shown below in black. **b** Percentage change in GABA concentration from the sham session to the active session plotted for all participants. Error bars show the SEM. **c** GABA/Glx ratio plotted for all participants for the sham and active sessions. Linear mixed effects modelling showing the was no significant effect of session (active or sham TUS). **d** Plot showing significant negative relationship between the percentage change in GABA concentration between sessions and the change in slope for pain intensity ratings in the active and sham conditions between T1 and T2 timepoints – representing the analgesic effect of TUS.

### 2.5 Safety profile of TUS stimulation

The stimulation was well tolerated by all participants. At the end of each session participants completed a symptom report questionnaire, which asked them to rate the intensity of a list of symptoms they may have experienced since the stimulation. After active TUS, participants reported the following symptoms: headache (n=2), scalp sensations (n=1), itchiness (n=1), double vision (n=1), twitching (n=2), balance problems (n=1), changes in hand movement (n=1), tingling sensations (n=4), muscle tightness (n=1), unusual feelings (n=1), anxiety (n=1), sleepiness (n=3), dizziness (n=3). All of these were reported as mild (defined as ‘present but not bothersome) except for one report of sleepiness which was reported as moderate (defined as ‘tolerable - required some intervention/medication but did not interfere with day-to-day activities’). After the sham session, participants reported: headache (n=3), neck pain (n=3), scalp sensations (n=1), itchiness (n=1), double vision (n=1), twitching (n=5), balance problems (n=1), changes in hand movement (n=2), tingling sensations (n=4), unusual feelings (n=1), anxiety (n=2), sleepiness (n=3), dizziness (n=1). All of these symptoms were reported as mild. No symptoms across either condition were reported as severe. Participants did note that tingling sensations and dizziness symptoms were likely due to the tonic cold stimulus and standing up after lying in the MRI scanner, respectively. Most of the reported symptoms were rated as unrelated or unlikely to be related to the stimulation.

## 3. Discussion

This study provides novel evidence that multi-focal TUS applied to dACC can modulate pain-related neural processing in healthy participants, even in the absence of significant immediate changes in subjective pain intensity. While no significant overall effect on pain ratings was observed between active and sham conditions at any single time point, there was a significantly greater reduction in pain ratings from ∼28 minutes to ∼55 minutes post-stimulation in the active TUS condition, suggesting a potential delayed analgesic effect. In parallel, altered sensory encoding was observed, with the typical relationship between cold temperature and pain intensity disrupted following active TUS. These behavioural findings were accompanied by significant changes in functional connectivity patterns, with increased coupling between the dACC and regions implicated in pain modulation and motor planning, including the supplementary motor area, pre-motor cortex, and anterior insular cortex, as well as decreased connectivity with PAG, a key hub in descending pain inhibition. Together, these findings suggest that TUS to the dACC can induce reorganisation within both cortical and subcortical pain networks, offering mechanistic insight into its potential utility as a non-invasive neuromodulatory intervention for pain treatment. The significant negative relationship between the percentage change in GABA concentration and the analgesic effect of TUS indicates that acute shifts in the neurochemistry balance within the dACC may underpin TUS effects, with participants having the largest decrease in GABA reporting the largest decrease in pain towards the last timepoint of the session.

Broadly, the findings of this study showing altered neural processing following TUS are aligned to previous literature demonstrating neuromodulatory effects of TUS applied to the dACC^4,20^. In addition, the finding that TUS to the dACC resulted in altered pain processing is also aligned to previous studies demonstrating modulation of pain responses^7,12^. More specifically, the two previous studies investigating TUS to the dACC for pain modulation had quite different study designs to our study, with the first investigating effects of online TUS on heat pain in healthy participants^7^ and the second investigating effects of offline TUS to reported pain in patients with chronic pain conditions^12^, making direct comparisons of the results challenging. Both studies did report significant reductions in pain ratings overall, which was not replicated in the current study.

The role of the dACC in pain processing has been well characterised in previous studies, with a key role in the affective component of pain responses, relating to aversiveness or unpleasantness associated with the pain stimulus^21-24^. The VAS for pain intensity used in many studies, including this one, relates primarily to the sensory-discriminative aspect of pain, but since the dACC is more involved in the emotional experience of pain, outcome measures such as pain unpleasantness, mood and emotional responses may have been more sensitive for assessment of TUS effects on this brain region^9^. This is reflected in the DBS literature, with multiple studies reporting that DBS to the dACC for chronic pain results in significant improvements in quality of life but not in VAS pain intensity ratings^10,25^, although longer term studies do show a significant reduction in numerical pain ratings at 6 months post-surgery^11,26^.

The greater decrease in active vs. sham conditions between T1 and T2 (potentially showing an analgesic effect at 55 minutes post-TUS) may indicate that TUS effects took longer to develop than anticipated, and if another later time point were included in the design (e.g. 1h or more post TUS), the TUS effects may have been stronger. Data from a macaque study conducted to investigate the temporal dynamics of TUS showed significantly altered functional connectivity between 43 minutes and 62 minutes post stimulation, with additional measures of intrinsic brain function showing effects up to 100 minutes (the last timepoint recorded)^27^. This is aligned to a study in healthy humans showing changes in functional connectivity shown across a larger number of regions at 46 minutes post-TUS compared to 13 minutes post-TUS, and significant changes in salience network and default mode network connectivity between sham and active TUS conditions at 46 minutes only (but not at 13 minutes)^4^. Overall, there are varying reports on the duration of TUS effects, with some studies reporting far longer-term effects, and wide variability in the measures used. For example, effects of TUS on reported pain intensity have been reported up to 7 days post-stimulation in chronic pain patients^12^, and *in vitro* research has shown that enhanced neuronal excitability in primary rat cortical neurons lasts up to 8 hours^28^. Further research is needed to understand the temporal dynamics of the effects of offline TUS, and to elucidate the neural mechanisms underpinning these effects across short-term (minutes), mid-term (hours) and long-term (days) timescales.

The study did have several limitations. The sample size is relatively small, with 29 participants for fMRI and 23 for MRS, limiting the statistical power of analyses and potentially impacting generalisability of the findings. In relation to the fMRI data, although a multi-echo acquisition was used, the final analysis was limited to data from a single echo time due to excessive noise in the combined data. As a result, we were unable to take advantage of the potential benefits of multi-echo processing, such as improved signal-to-noise ratio. Furthermore, the selected echo time was not specifically optimised for use in our analysis, which may have affected sensitivity. However, given the substantial noise in the combined data, using the second echo alone provided a more reliable and interpretable dataset. It is also worth noting that the scanner software update completed in the middle of data collection necessitated a change in the scan sequence, which may have introduced additional variability in data quality.

A strength of the study was the double-blinded design, which aimed to reduce any bias arising from both participants and researchers towards active vs. sham stimulation. Blinding effectiveness was assessed for the researcher administering the stimulation in a sample of sessions (n=34), with exactly 50% accuracy rate, indicating the blinding procedures in place for researchers were very effective. For participants, the sound delivered via bone conducting headphones during the sham session aimed to closely match the active TUS, but blinding effectiveness was not assessed. Assessment of blinding effectiveness for participants would have been valuable to evaluate any bias potentially introduced by participant’s expectations. The researcher conducting data analysis was unblinded at that stage, which could also be considered a limitation of the study as it could have introduced bias.

Overall, this study provides novel mechanistic insight into how multi-focal TUS targeting the dACC can alter pain-related brain function in healthy individuals. Together, these findings demonstrate the capacity of TUS to influence distributed brain circuits involved in the complex experience of pain, supporting its potential as a targeted, non-invasive intervention. These results highlight the importance of including longer post-stimulation assessment periods in future studies, as well as expanding outcome measures to better capture the affective and evaluative dimensions of pain processing.

In conclusion, this study extends current understanding of the effects of TUS on the dACC by demonstrating functionally meaningful changes in pain perception and brain connectivity. The findings lay important groundwork for future translational research exploring the therapeutic use of TUS in chronic pain populations. Further work in larger samples, using multimodal approaches and longer follow-up periods, will be critical to fully characterise the efficacy, duration, and mechanisms of TUS-induced modulation of pain.

## 4. Methods

### 4.1 Participants and approvals

The study was approved by the University of Plymouth Faculty of Health Staff Research Ethics and Integrity Committee (reference: 4183). Exclusion criteria were applied for TUS and MRI safety and to ensure participants were healthy, with no pain-related conditions. All participants gave written informed consent prior to taking part.

### 4.2 Study design

The study design was a double-blind, randomised cross-over study in healthy participants. The study consisted of 3 visits: the first to acquire T1-weighted and PETRA images for acoustic simulations, followed by two main TUS-fMRI sessions with either active TUS or sham (Fig. 1A). At each main session, first active/sham TUS was applied during a tonic cold stimulus, then an MRI scan was completed with fMRI blocks during baseline, tonic cold and recovery, and a magnetic resonance spectroscopy (MRS) block during tonic cold.

### 4.3 Tonic pain model

Tonic pain was induced using a modified cold pressor test^29,30^ with gelled water (temp range: 3.9±1.8oC). The gelled water was produced as described by Lapotka et al., using water, a thickening product containing modified cornstarch, and salt^31^. Verbal pain intensity ratings were collected after each of the three tonic cold pain stimuli, on a numerical rating scale (NRS) from 0

– no pain, to 100 – worst pain imaginable.

### 4.4 TUS stimulation protocol

The NeuroFUS transducer power output (TPO) system and CTX-500-4 transducer *(Brainbox Ltd*., *Cardiff, UK)*; a four-element annular transducer (diameter = 64 mm, central frequency = 500 kHz, with steering range between 27.3 and 82.6mm), was used to deliver the TUS protocol. The dACC targeting and optimal transducer placement was individually planned using structural MRI scans (T1-weighted and PETRA) and k-Plan software *(BrainBox, Inc.)* to run acoustic and thermal simulations for each participant, to ensure accurate pressure delivery at the target area and adherence to safety protocols. The three dACC targets were defined anatomically using MNI co-ordinates: target A: -3,37,12; target B: -3,33,17; and target C: -3,28,22. Repetitive TUS followed a 10Hz-patterned protocol with a 10% duty cycle (fundamental frequency = 500KHz, pulse duration = 10ms, pulsed every 100ms), applied for 80 seconds at each target sequentially (4 minutes in total). The I_SPPA_ in water was maintained at 54W/cm^2^ across participants. We adhered to safety guidelines for human ultrasound exposure as defined by the International Thermal and Radiological Ultrasound Safety Standards and Thresholds (ITRUSST ^19^).

To ensure ultrasound transmission, a layer of ultrasound gel *(Aquasonic 100, Parker Laboratories Inc.)* was applied at the transducer placement site, with a 2cm gel pad *(Aquaflex, Parker Laboratories Inc.)* positioned between the transducer and the participant’s head. Hair was not shaved; while applying the layer of gel the hair was carefully combed and smoothed to eliminate air gaps. Neuronavigation was performed using Brainsight v2.5 *(Rogue Research Inc*., *Montréal, Québec, Canada)* with T1-weighted anatomical MR images. During each session, focal depth measurements to each target obtained from Brainsight were entered into the NeuroFUS TPO system before stimulation, and the trajectory was sampled to support post-session confirmatory acoustic simulations.

The sham condition consisted of a matched auditory stimulus delivered to participants via bone conducted headphones. Since the study was double-blind, all procedures were matched at both sessions (with the same neuro-navigation set-up and transducer placement at each target). An independent researcher managed the blinding and administered either active or sham TUS depending on the randomisation for each participant. Data was unblinded prior to analysis.

### 4.5 Imaging data acquisition (MRI and MRS)

Data were collected on a Siemens MAGNETOM Prisma 3T scanner with a 64-channel head coil. At visit 1, a T1-weighted structural scan and a pointwise encoding time reduction with radial acquisition (PETRA) scan were collected for use in acoustic simulations and neuronavigation. The T1-weighted structural scan was acquired with an MPRAGE sequence with a TR of 2.1s, echo time of 2.26ms, inversion time of 900ms, flip angle of 8°, GRAPPA acceleration factor of 2, matrix size of 256×256, 176 slices, and with 1mm^3^ isotropic voxels). The PETRA was acquired with TR of 3.61ms, echo time of 0.07ms, flip angle of 8°, 320 slices per slab and slice thickness of 0.75mm. 3D distortion correction was applied following acquisition. The PETRA scan was converted to a pseudo-CT scan for use in acoustic simulations using the PETRA-to-CT MATLAB toolbox^32^.

At visits 2 and 3, three resting-state BOLD scans were completed (rest/baseline, tonic cold pain, and post-pain recovery), followed by a field map and T1-weighted structural scan, and finally a magnetic resonance spectroscopy (MRS) scan. Due to an upgrade of the scanner software, the second 14 datasets were collected with a slightly modified scan sequence than the first 15 (for BOLD and MRS scans only). Modified parameters are indicated in italic text following the initial parameter used if applicable. Resting-state BOLD fMRI data were acquired using a multi-echo echo-planar imaging (EPI) sequence with repetition time (TR) of 1.5s */ 1.55s*, 4 echo times (11.0, 27.25, 43.5, and 59.75ms */ 13.6, 29.86, 46.1 and 62.36ms*), flip angle of 77°, 2.6mm^3^ isotropic voxels, 51 axial slices, interleaved acquisition, 220mm field of view, matrix size of 84×84, 2.6mm slice thickness, GRAPPA acceleration factor of 2, multi-band acceleration factor of 3, bandwidth of 2480Hz/Px, and 240 volumes per run. The field map was acquired to enable correction of field inhomogeneity during analysis, with 2.6mm^3^ isotropic voxels and 220mm field of view. The T1-weighted scan had the same parameters as for visit 1. MRS data was acquired in a 3.5×2.5×2.5cm^3^ voxel positioned in the dACC (TUS target region) using a MEGA-PRESS sequence to quantify GABA concentrations, with TR of 2s, echo time of 68ms, excitation flip angle of 90°, and 256 */ 128* averages. Water suppression was achieved using a frequency-selective saturation pulse (bandwidth = 35Hz) */ VAPOUR (bandwidth = 60Hz)*, and MEGA editing pulses were centred at 1.9ppm and 7.5ppm to detect GABA, with editing pulse flip angle of 180°. Spectral data were collected with a vector size of 2048, bandwidth of 2000Hz */ 1850Hz*, and acquisition duration of 1024ms */ 1107ms*. A water unsuppressed reference was collected with 8 averages. Automatic frequency and shim adjustments were applied prior to acquisition.

### 4.6 Post-TUS symptom questionnaire

After each session participants were asked to complete a symptom questionnaire, which consisted of a list of symptoms for which they were asked to rate the intensity they experienced the symptom using a 4-point scale (absent, mild, moderate, severe) and whether they thought their experience was related to the stimulation on a 5-point scale (unrelated, unlikely, possible, probable, definite). Participants could note up to two additional symptoms and could provide comments. See Supplementary Figure 1 for the full questionnaire.

### 4.7 Statistical analysis

For behavioural data, the relationship between pain intensity ratings and TUS condition (as well as additional potentially confounding factors) were assessed using linear mixed effects modelling in R v4.4.0. The following general liner model (GLM) was used:

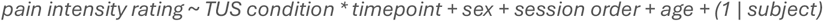

The relationship between pain intensity rating and temperature of the tonic cold stimulus in both the active and sham TUS sessions was assessed using simple linear regression in GraphPad PRISM v10.1.0.

Resting state fMRI data were analysed using tools in FMRIB Software Library v6.0 (FSL)^33-35^. Structural and magnitude images were brain extracted^36^ and a calibrated field map image was prepared as required for B0 unwarping. Due to excessive noise in the multi-echo data, only the second echo time (TE2; 27.25 / *29.86*) was used for analysis of the functional data, and for this report we focussed only on the tonic pain block. Pre-processing steps including registration to structural and MNI standard images, B0 unwarping, motion correction, spatial smoothing (5mm) and high-pass temporal filtering, conducted using FEAT^37^. For seed-based analysis, the time course of BOLD activity from the dACC seed-region was extracted and used to generate individual statistical maps of the functionally correlated activity across the whole brain for each participant, using the GLM approach implemented with FEAT^37^. Motion outliers identified using the fslmotionoutliers tool, average signal from the white matter and cerebrospinal fluid generated using anatomical segmentations for each tissue type, and the six motion parameters from the motion correction step were included as nuisance covariates. Group-level whole brain, mixed effects analysis with a cluster-based correction for multiple comparisons was performed using FEAT to search for differences in functional connectivity between the active TUS and sham TUS conditions^38^. The mask for the seed-region was generated by created 5mm spheres around each of the 3 targets, combining these, and then warping this mask to each individual participant’s functional space. For each participant, any regions of the dACC mask that overlapped with thresholded white matter or cerebrospinal fluid masks were subtracted, to create an individualised seed in grey matter only. To explore changes in resting state network connectivity, probabilistic independent component analysis as implemented in MELODIC was used^39-41^. Data from both active and sham TUS sessions were pre-processed by masking non-brain voxels, applying voxel-wise demeaning and normalisation of the voxel-wise variance, then projected into a 20-dimensional subspace using principal component analysis. Spatial maps from this group-average analysis were used to generate subject-specific versions, and associated timeseries, using dual regression^42,43^. Group differences between the active TUS and sham TUS conditions were then compared using randomise with threshold-free cluster enhancement (TFCE; 5000 permutations, p<0.05)^44^.

For MRS, data were analysed using the MATLAB-based Gannet 3.4.0 toolkit^45^. Processing steps included frequency and phase correction conducted with robust spectral registration^46^, 3Hz exponential line broadening, and individual transient averaging. The GABA and Glx peaks at 3.0ppm and 3.75ppm, respectively, were modelled relative to water and corrected for voxel tissue composition using segmentation of T1-weighted structural images conducted with SPM^13,47^. Data quality was assessed via visual inspection for spectral artefacts and on quality metrics including FWHM, GABA+ signal to noise ratio (SNR) and model fit error, resulting in 3 datasets being excluded from analysis. One further dataset was excluded as an outlier, identified using the ROUT method (Q=1%) in GraphPad PRISM, resulting in 23 datasets included for analysis. The relationship between metabolite concentrations (GABA, GLx and GABA/Glx ratio) and TUS condition (as well as additional potentially confounding factors) were assessed using linear mixed effects modelling in R v4.4.0. The following GLM was used for this:

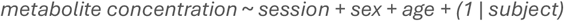

The relationship between the percentage change in GABA concentration between sessions and the change in slope for pain intensity ratings in the active and sham conditions between T1 and T2 timepoints was assessed using simple linear regression in GraphPad PRISM v10.1.0.

## 5. Data availability

Data supporting the findings of this study are available at https://osf.io/vp2gz/.

## Acknowledgements

We thank Jamie Roberts for supporting the development of the scanning protocols used in the study. Thank you to staff at the Brain Research and Imaging Centre (BRIC) for support with scanning, and to all our participants for taking part in the research.

Sam Hughes, Sophie Clarke and Elsa Fouragnan are funded by a Neuromod+ grant (EP/W035057/1). Elsa Fouragnan is funded by a UKRI FLF (MR/Y034368/1), a BBSRC (BB/Y001494/1) and an ARIA grant (SCNI-PR01-P15).

## 8. Author Contributions Statement

These authors contributed equally: Elsa Fouragnan, Sam Hughes

### Contributions

Conceptualization: SC, EF and SH. Data curation: SC, ML, NB, EF. Formal analysis: SC. Funding acquisition: EF and SH. Investigation: SC, SM and JM. Methodology: SC, ML, NB, EF and SH. Project administration: SC, SM and JM. Resources: EF and SH. Software: ML and NB. Supervision: EF and SH. Validation: SC, SM, ML, JM, NB, EF and SH. Visualization: SC and EF. Writing – original draft: SC. Writing – review & editing: SC, SM, ML, JM, NB, EF and SH.

## References

1 Bault, N., Yaakub, S. N. & Fouragnan, E. Early-phase neuroplasticity induced by offline transcranial ultrasound stimulation in primates. Current Opinion in Behavioral Sciences 56, 101370 (2024). 10.1016/j.cobeha.2024.101370

2 Murphy, K. & Fouragnan, E. The future of transcranial ultrasound as a precision brain interface. PLoS Biol 22, e3002884 (2024). 10.1371/journal.pbio.3002884

3 Pellow, C., Pichardo, S. & Pike, G. B. A systematic review of preclinical and clinical transcranial ultrasound neuromodulation and opportunities for functional connectomics. Brain Stimul 17, 734–751 (2024). 10.1016/j.brs.2024.06.005

4 Yaakub, S. N. et al. Transcranial focused ultrasound-mediated neurochemical and functional connectivity changes in deep cortical regions in humans. Nat Commun 14, 5318 (2023). 10.1038/s41467-023-40998-0

5 Badran, B. W. & Peng, X. Transcranial focused ultrasound (tFUS): a promising noninvasive deep brain stimulation approach for pain. Neuropsychopharmacology 49, 351–352 (2024). 10.1038/s41386-023-01699-w

6 Badran, B. W. et al. Sonication of the anterior thalamus with MRI-Guided transcranial focused ultrasound (tFUS) alters pain thresholds in healthy adults: A double-blind, sham-controlled study. Brain Stimul 13, 1805–1812 (2020). 10.1016/j.brs.2020.10.007

7 Strohman, A., Payne, B., In, A., Stebbins, K. & Legon, W. Low-Intensity Focused Ultrasound to the Human Dorsal Anterior Cingulate Attenuates Acute Pain Perception and Autonomic Responses. J Neurosci 44 (2024). 10.1523/jneurosci.1011-23.2023

8 Hameroff, S. et al. Transcranial Ultrasound (TUS) Effects on Mental States: A Pilot Study. Brain Stimulation 6, 409–415 (2013). 10.1016/j.brs.2012.05.002

9 Russo, J. F. & Sheth, S. A. Deep brain stimulation of the dorsal anterior cingulate cortex for the treatment of chronic neuropathic pain. Neurosurg Focus 38, E11 (2015). 10.3171/2015.3.Focus1543

10 Boccard, S. G. et al. Targeting the affective component of chronic pain: a case series of deep brain stimulation of the anterior cingulate cortex. Neurosurgery 74, 628-635; discussion 635-627 (2014). 10.1227/neu.0000000000000321

11 Boccard, S. G. J. et al. Long-Term Results of Deep Brain Stimulation of the Anterior Cingulate Cortex for Neuropathic Pain. World Neurosurg 106, 625–637 (2017). 10.1016/j.wneu.2017.06.173

12 Riis, T. S., Feldman, D. A., Losser, A. J., Okifuji, A. & Kubanek, J. Noninvasive targeted modulation of pain circuits with focused ultrasonic waves. Pain 165, 2829–2839 (2024). 10.1097/j.pain.0000000000003322

13 Ashburner, J. & Friston, K. J. Unified segmentation. Neuroimage 26, 839–851 (2005). 10.1016/j.neuroimage.2005.02.018

14 Chou, T. et al. Transcranial focused ultrasound of the amygdala modulates fear network activation and connectivity. Brain Stimulation 17, 312–320 (2024). 10.1016/j.brs.2024.03.004

15 Kuhn, T. et al. Transcranial focused ultrasound selectively increases perfusion and modulates functional connectivity of deep brain regions in humans. Frontiers in Neural Circuits Volume 17 - 2023 (2023). 10.3389/fncir.2023.1120410

16 Lord, B. et al. Transcranial focused ultrasound to the posterior cingulate cortex modulates default mode network and subjective experience: an fMRI pilot study. Frontiers in Human Neuroscience Volume 18 - 2024 (2024). 10.3389/fnhum.2024.1392199

17 Jung, J., Atkinson-Clement, C., Kaiser, M. & Ralph, M. A. L. Transcranial focused ultrasound stimulation of the anterior temporal lobe enhances semantic memory by modulating brain morphology, neurochemistry and neural dynamics. bioRxiv, 2025.2003.2010.642483 (2025). 10.1101/2025.03.10.642483

18 Yang, P. S. et al. Transcranial focused ultrasound to the thalamus is associated with reduced extracellular GABA levels in rats. Neuropsychobiology 65, 153–160 (2012). 10.1159/000336001

19 Aubry, J.-F. et al. ITRUSST consensus on biophysical safety for transcranial ultrasonic stimulation. arXiv preprint 2311.05359 (2023).

20 Koutsoumpari, N. et al. Ultrasound neuromodulation reveals distinct roles of the dorsal anterior cingulate cortex and anterior insula in Pavlovian biases. bioRxiv, 2025.2006.2012.659273 (2025). 10.1101/2025.06.12.659273

21 Fuchs, P. N., Peng, Y. B., Boyette-Davis, J. A. & Uhelski, M. L. The anterior cingulate cortex and pain processing. Front Integr Neurosci 8, 35 (2014). 10.3389/fnint.2014.00035

22 Lieberman, M. D. & Eisenberger, N. I. The dorsal anterior cingulate cortex is selective for pain: Results from large-scale reverse inference. Proceedings of the National Academy of Sciences 112, 15250–15255 (2015). 10.1073/pnas.1515083112

23 Rainville, P., Duncan, G. H., Price, D. D., Carrier, B.t. & Bushnell, M. C. Pain Affect Encoded in Human Anterior Cingulate But Not Somatosensory Cortex. Science 277, 968–971 (1997). 10.1126/science.277.5328.968

24 Singh, A. et al. Mapping Cortical Integration of Sensory and Affective Pain Pathways. Current Biology 30, 1703-1715.e1705 (2020). 10.1016/j.cub.2020.02.091

25 Fontaine, D. et al. Safety and feasibility of deep brain stimulation of the anterior cingulate and thalamus in chronic refractory neuropathic pain: a pilot and randomized study. The Journal of Headache and Pain 26, 35 (2025). 10.1186/s10194-025-01967-8

26 Levi, V. et al. Dorsal anterior cingulate cortex (ACC) deep brain stimulation (DBS): a promising surgical option for the treatment of refractory thalamic pain syndrome (TPS). Acta Neurochir (Wien) 161, 1579–1588 (2019). 10.1007/s00701-019-03975-5

27 Atkinson-Clement, C. et al. Temporal dynamics of offline transcranial ultrasound stimulation. Current Research in Neurobiology 8, 100148 (2025). 10.1016/j.crneur.2025.100148

28 Clennell, B. et al. Transient ultrasound stimulation has lasting effects on neuronal excitability. Brain Stimul 14, 217–225 (2021). 10.1016/j.brs.2021.01.003

29 Medina, S. & Hughes, S. W. A Novel Investigation of an In-Scanner Alternative to the Cold Pressor Test in Healthy Individuals. Hum Brain Mapp 46, e70291 (2025). 10.1002/hbm.70291

30 Medina, S. & Hughes, S. W. Immersion in nature through virtual reality attenuates the development and spread of mechanical secondary hyperalgesia: a role for insulo-thalamic effective connectivity. Pain (2025). 10.1097/j.pain.0000000000003701

31 Lapotka, M., Ruz, M., Salamanca Ballesteros, A. & Ocón Hernández, O. Cold pressor gel test: A safe alternative to the cold pressor test in fMRI. Magnetic Resonance in Medicine 78, 1464–1468 (2017). 10.1002/mrm.26529

32 Treeby, B. PETRA-TO-CT, <https://github.com/ucl-bug/petra-to-ct (2023).

33 Jenkinson, M., Beckmann, C. F., Behrens, T. E., Woolrich, M. W. & Smith, S. M. FSL. Neuroimage 62, 782–790 (2012). 10.1016/j.neuroimage.2011.09.015

34 Smith, S. M. et al. Advances in functional and structural MR image analysis and implementation as FSL. Neuroimage 23 Suppl 1, S208–219 (2004). 10.1016/j.neuroimage.2004.07.051

35 Woolrich, M. W. et al. Bayesian analysis of neuroimaging data in FSL. Neuroimage 45, S173–186 (2009). 10.1016/j.neuroimage.2008.10.055

36 Smith, S. M. Fast robust automated brain extraction. Hum Brain Mapp 17, 143–155 (2002). 10.1002/hbm.10062

37 Woolrich, M. W., Ripley, B. D., Brady, M. & Smith, S. M. Temporal autocorrelation in univariate linear modeling of FMRI data. Neuroimage 14, 1370–1386 (2001). 10.1006/nimg.2001.0931

38 Woolrich, M. W., Behrens, T. E., Beckmann, C. F., Jenkinson, M. & Smith, S. M. Multilevel linear modelling for FMRI group analysis using Bayesian inference. Neuroimage 21, 1732–1747 (2004). 10.1016/j.neuroimage.2003.12.023

39 Beckmann, C. F. & Smith, S. M. Probabilistic independent component analysis for functional magnetic resonance imaging. IEEE Transactions on Medical Imaging 23, 137–152 (2004). 10.1109/TMI.2003.822821

40 Beckmann, C. F. & Smith, S. M. Tensorial extensions of independent component analysis for multisubject FMRI analysis. Neuroimage 25, 294–311 (2005). 10.1016/j.neuroimage.2004.10.043

41 Hyvarinen, A. Fast and robust fixed-point algorithms for independent component analysis. IEEE Transactions on Neural Networks 10, 626–634 (1999). 10.1109/72.761722

42 Beckmann, C. F., Mackay, C. E., Filippini, N. & Smith, S. M. Group comparison of resting-state FMRI data using multi-subject ICA and dual regression. NeuroImage 47, S148 (2009). 10.1016/S1053-8119(09)71511-3

43 Nickerson, L. D., Smith, S. M., Öngür, D. & Beckmann, C. F. Using Dual Regression to Investigate Network Shape and Amplitude in Functional Connectivity Analyses. Front Neurosci 11, 115 (2017). 10.3389/fnins.2017.00115

44 Winkler, A. M., Ridgway, G. R., Webster, M. A., Smith, S. M. & Nichols, T. E. Permutation inference for the general linear model. Neuroimage 92, 381–397 (2014). 10.1016/j.neuroimage.2014.01.060

45 Edden, R. A. E., Puts, N. A. J., Harris, A. D., Barker, P. B. & Evans, C. J. Gannet: A batch-processing tool for the quantitative analysis of gamma-aminobutyric acid–edited MR spectroscopy spectra. Journal of Magnetic Resonance Imaging 40, 1445–1452 (2014). 10.1002/jmri.24478

46 Mikkelsen, M. et al. Correcting frequency and phase offsets in MRS data using robust spectral registration. NMR in Biomedicine 33, e4368 (2020). 10.1002/nbm.4368

47 Harris, A. D., Puts, N. A. J. & Edden, R. A. E. Tissue correction for GABA-edited MRS: Considerations of voxel composition, tissue segmentation, and tissue relaxations. Journal of Magnetic Resonance Imaging 42, 1431–1440 (2015). 10.1002/jmri.24903

