## Supplementary Table 1 for "Multi-focal ultrasound neuromodulation to the dorsal anterior cingulate cortex disrupts behavioural and neural pain processing"

Supplementary Table 1: Individual post-stimulation simulation results

| Participant | Intensity (I_SPPA_) in dACC | | | Pressure (kPa) in dACC | | | Thermal dose in soft tissues (CEM43^o^C) | | | Mechanical index in soft tissues (MI) | | |
| --- | --- | --- | --- | --- | --- | --- | --- | --- | --- | --- | --- | --- |
|  | A | B | C | A | B | C | A | B | C | A | B | C |
| 1 | 2.85 | 2.09 | 4.36 | 324.89 | 269.74 | 391.89 | 0.001 | 0.002 | 0.003 | 0.40 | 0.36 | 0.50 |
| 2 | 2.83 | 2.96 | 3.51 | 301.01 | 306.88 | 352.95 | 0.001 | 0.002 | 0.003 | 0.40 | 0.42 | 0.44 |
| 3 | 1.51 | 2.41 | 5.29 | 230.88 | 289.45 | 443.00 | 0.003 | 0.005 | 0.010 | 0.73 | 0.87 | 0.80 |
| 4 | 2.88 | 3.08 | 3.94 | 325.93 | 336.46 | 364.07 | 0.004 | 0.009 | 0.012 | 0.86 | 0.92 | 1.56 |
| 5 | 4.18 | 5.40 | 5.66 | 392.47 | 437.13 | 455.67 | 0.005 | 0.010 | 0.015 | 1.00 | 0.79 | 0.86 |
| 6 | 1.77 | 5.44 | 6.32 | 257.96 | 433.93 | 475.64 | 0.002 | 0.003 | 0.006 | 0.70 | 0.70 | 0.71 |
| 7 | 7.04 | 6.11 | 5.61 | 495.60 | 466.31 | 448.07 | 0.003 | 0.004 | 0.008 | 0.83 | 0.75 | 0.86 |
| 8 | 3.27 | 2.96 | 5.27 | 347.82 | 318.48 | 419.19 | 0.003 | 0.009 | 0.012 | 0.80 | 0.74 | 0.70 |
| 9 | 4.61 | 5.31 | 5.42 | 417.72 | 453.10 | 440.30 | 0.002 | 0.004 | 0.006 | 0.57 | 0.65 | 0.77 |
| 10 | 5.28 | 5.29 | 5.05 | 434.98 | 434.41 | 408.89 | 0.002 | 0.004 | 0.007 | 0.77 | 0.70 | 0.73 |
| 11 | 2.47 | 2.45 | 3.24 | 297.54 | 280.90 | 348.51 | 0.005 | 0.009 | 0.015 | 0.69 | 1.07 | 1.02 |
| 12 | 3.07 | 5.01 | 5.25 | 328.56 | 409.87 | 438.71 | 0.002 | 0.004 | 0.005 | 0.74 | 0.75 | 0.86 |
| 13 | 2.05 | 4.18 | 6.03 | 275.37 | 370.41 | 460.00 | 0.002 | 0.004 | 0.007 | 0.66 | 0.55 | 0.59 |
| 14 | 0.52 | 0.54 | 5.56 | 138.29 | 140.23 | 436.37 | 0.003 | 0.009 | 0.012 | 0.63 | 0.65 | 0.99 |
| 15 | 5.89 | 5.39 | 5.11 | 452.77 | 440.09 | 428.83 | 0.002 | 0.007 | 0.009 | 0.72 | 0.71 | 0.86 |
| 16 | 3.05 | 4.60 | 5.81 | 324.19 | 400.38 | 455.95 | 0.002 | 0.006 | 0.013 | 0.67 | 0.70 | 1.00 |
| 17 | 1.61 | 2.16 | 3.24 | 245.08 | 281.19 | 325.95 | 0.002 | 0.005 | 0.008 | 0.56 | 0.63 | 0.66 |
| 18 | 1.39 | 1.39 | 1.93 | 222.26 | 227.24 | 263.20 | 0.002 | 0.007 | 0.012 | 0.67 | 0.82 | 1.00 |
| 19 | 2.36 | 2.05 | 3.55 | 290.67 | 271.99 | 358.04 | 0.002 | 0.006 | 0.012 | 0.73 | 0.77 | 1.15 |
| 20 | 0.95 | 1.55 | 2.14 | 182.20 | 238.80 | 271.71 | 0.002 | 0.007 | 0.009 | 0.74 | 0.79 | 0.84 |
| 21 | 7.73 | 6.87 | 4.57 | 513.32 | 481.16 | 407.82 | 0.002 | 0.009 | 0.016 | 0.86 | 0.99 | 1.14 |
| 22 | 2.06 | 2.34 | 3.21 | 263.06 | 288.15 | 330.84 | 0.003 | 0.011 | 0.021 | 0.75 | 1.00 | 1.08 |
| 23 | 2.89 | 3.11 | 2.45 | 327.72 | 331.35 | 292.29 | 0.003 | 0.007 | 0.011 | 0.70 | 0.64 | 0.84 |
| 24 | 2.96 | 2.91 | 2.71 | 328.76 | 319.75 | 315.04 | 0.002 | 0.006 | 0.010 | 0.78 | 0.63 | 0.75 |
| 25 | 4.06 | 5.08 | 3.50 | 381.40 | 427.61 | 351.94 | 0.002 | 0.006 | 0.011 | 0.63 | 0.70 | 0.69 |
| 26 | 4.69 | 3.69 | 4.53 | 404.10 | 344.64 | 419.09 | 0.003 | 0.013 | 0.023 | 0.89 | 0.97 | 0.76 |
| 27 | 5.16 | 5.22 | 7.27 | 433.54 | 408.61 | 500.96 | 0.002 | 0.006 | 0.012 | 0.72 | 0.74 | 1.15 |
| 28 | 2.56 | 2.43 | 1.98 | 304.72 | 296.65 | 266.94 | 0.002 | 0.006 | 0.010 | 0.65 | 0.83 | 0.70 |
| 29 | 2.80 | 2.86 | 3.32 | 323.47 | 316.43 | 352.58 | 0.002 | 0.006 | 0.012 | 0.55 | 0.61 | 0.73 |
| 30 | 3.23 | 3.88 | 3.79 | 348.91 | 354.64 | 363.48 | 0.003 | 0.026 | 0.036 | 0.90 | 1.14 | 1.04 |
| 31 | 4.73 | 6.74 | 6.71 | 402.30 | 477.72 | 496.89 | 0.002 | 0.005 | 0.010 | 0.63 | 0.78 | 0.73 |
| 32 | 1.53 | 1.47 | 2.50 | 236.99 | 223.78 | 298.74 | 0.002 | 0.006 | 0.012 | 0.92 | 0.78 | 0.95 |
