## Supplementary figures and images for "Multi-focal ultrasound neuromodulation to the dorsal anterior cingulate cortex disrupts behavioural and neural pain processing"

### Supplementary Figure 1

Supplementary Figure 1: Post-TUS Symptom Report Questionnaire

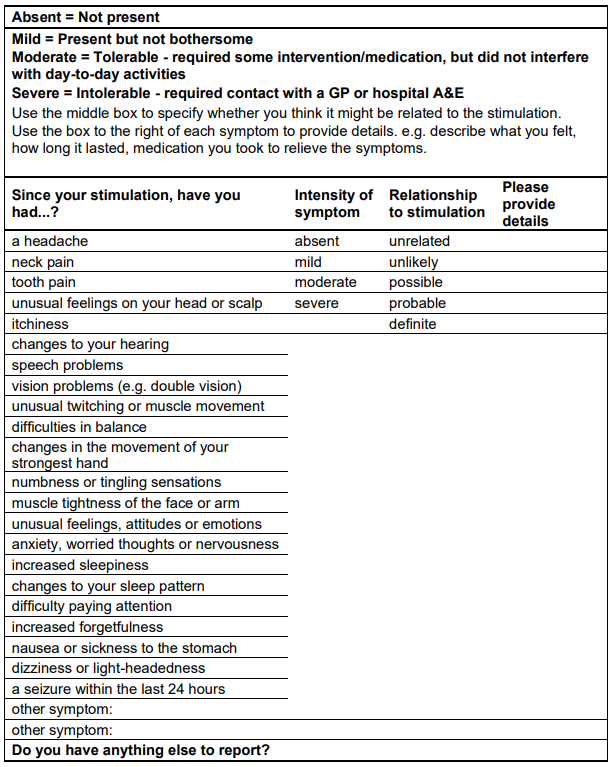
